# “Ancestralization” of human pluripotent stem cells by multiplexed precise genome editing

**DOI:** 10.1101/342311

**Authors:** Stephan Riesenberg, Tomislav Maricic, Svante Pääbo

**Affiliations:** Department of Evolutionary Genetics, Max Planck Institute for Evolutionary Anthropology

## Abstract

We show that inactivation of the DNA-dependent protein kinase catalytic subunit (DNA-PKcs) results in a drastic increase in efficiency of precise genome editing with CRISPR enzymes in human stem cells, allowing up to 79% of chromosomes to carry an intended nucleotide substitution when a single genomic site is targeted. When three different genes are simultaneously targeted, 12% of the isolated cells carry the targeted amino acid-changing substitutions in homozygous forms. These substitutions represent the first step towards resurrecting the proteome ancestral to Neandertals and modern humans. DNA-PKcs inactivation will greatly facilitate multiplexed precise genome editing in animal cells.

---

Genome sequences of our closest evolutionary relatives, Neandertals and Denisovans, allow the identification of novel genetic changes that are present in all or almost all humans today. These changes are of interest because some of them may be involved in phenotypes that made the cultural and technological developments associated with modern humans possible ^1^. However, because the ancestral versions of such changes are absent or extremely rare in present-day humans, their physiological consequences cannot be easily studied, especially not in a homozygous form. Furthermore, it is likely that combinations of such mutations may be necessary to induce relevant phenotypes. One way to study such changes is to introduce the ancestral genetic variants in human induced pluripotent stem cells (iPSCs) which can then be differentiated into various cell types ^2^ or used to generate organoids ^3,4^. Unfortunately, this approach is limited by the inability to introduce multiple precise genetic changes in mammalian cells. To overcome this limitation, we developed an approach to simultaneously introduce multiple nucleotide substitutions in a homozygous form in human stem cells.

Genome editing uses clustered regularly interspaced short palindromic repeats (CRISPR) nucleases, or other nucleases ^5^, to introduce targeted double-strand breaks (DSBs) in a genome. These are then repaired by two competing cellular pathways: non-homologous end-joining (NHEJ) or homology-directed repair (HDR). Because NHEJ often introduces deletions or insertions (indels) it is often used to inactivate genes. In contrast, HDR utilizes a sister chromatid to repair the chromosome carrying a DNA break. This activity can be used for precise genome editing (PGE) if exogenous DNA carrying desired mutations is provided as a donor. However, NHEJ is typically more efficient than HDR, leading to low PGE efficiencies. As a result, multiplexed PGE has not been reported in animal cells while multiplexed introduction of insertions and deletions has been achieved. For example, eight alleles of five different genes have been inactivated in mouse embryonic stem cells by insertions or deletions ^6^ and in a porcine immortalized cell line 62 copies of an endogenous retrovirus have been inactivated by the use of a two guide RNAs ^7^. This shows that#DNA cleavage by CRISPR enzymes is not the rate-limiting step for genome editing. Rather, the higher efficiency of NHEJ relative to HDR limits the ability to introduce precise substitutions. Several studies have therefore tried to inhibit NHEJ in order to increase the efficiencies of PGE, for example by blocking the synthesis of NHEJ proteins with siRNAs, by the expression of adenovirus proteins that degrade proteins necessary for NHEJ ^8^, and by using small molecules that block NHEJ ^9,10^. Others have tried to enhance HDR by cell cycle synchronization ^11,12^ and improved DNA donor design ^13-15^. A third approach that does not rely on HDR to introduce nucleotide substitutions is to use proteins that chemically induce nucleotide transitions but this allows only four out of 12 possible substitutions to be introduced ^16,17^.

We focused our efforts to increase HDR and thus PGE on the DNA-dependent protein kinase (DNA-PK) complex, which consists of the subunits Ku70, Ku80 and the DNA-PK catalytic subunit (DNA-PKcs) and covers the DNA ends after a DSB is introduced (Fig. 1A) ^18^. When bound to DNA ends, the DNA-PKcs undergoes autophosphorylation and recruits other proteins such as DNA ligase IV, XRCC4, and XLF ^18^ which are necessary for canonical NHEJ ^19,20^. If canonical NHEJ is compromised, a slower DNA-PKcs-independent alternative NHEJ pathway may take over ^21^. Alternatively, HDR can be induced if a protein complex composed of Mre11, Rad50 and Nbs1 binds to the DNA ends. This results in the generation of 3’ overhanging DNA ends, RAD51 nucleoprotein filament formation, and annealing of donor DNA that serve as template for DNA synthesis leading to the faithful repair of the DSB ^18,22^.

**Fig. 1:**
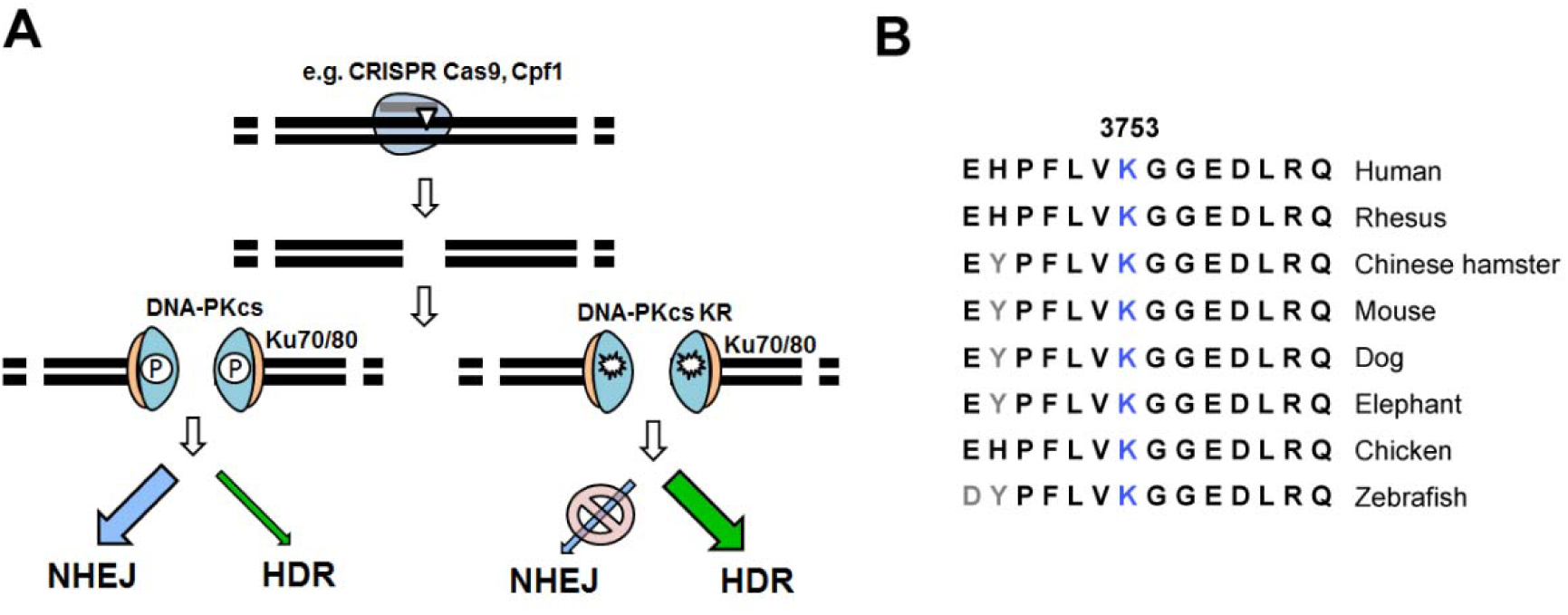
Catalytically inactive DNA-PKcs (K3753R) promotes HDR and blocks NHEJ. **(A)** After a double strand break (DSB) induced by *e.g*. CRISPR-Cas9 or Cpf1 (light blue with gray gRNA), DNA ends are covered by Ku70/80 (orange), followed by binding of DNA-PKcs (cyan blue), both constituting a DNA-PK complex. Autophosphorylation of DNA-PKcs leads to recruitment and activation of downstream NHEJ proteins. If DNA-PKcs is catalytically inactivated (e.g. by the K3753R mutation), autophosphorylation cannot take place and the NHEJ pathway is be blocked ^22,29^ leading to preferential HDR repair of the DSB. **(B)** The amino acid sequence of DNA-PKcs around K3753 is conserved among vertebrates (C) ^33^.

When bound to DSBs, the ~470kDa DNA-PKcs protein undergoes conformational changes induced at least in part by autophosphorylation ^23,24^ as well as by other related kinases *^25,26^* at more than 60 phosphorylation sites ^20,27,28^. When phosphorylated, some sites activate the kinase activity of DNA-PKcs, while others disengage c-NHEJ and inactive the kinase activity of DNA-PKcs (Fig. 1B). In 2008, it was shown that a lysine to arginine mutation at position 3753 near the ATP binding site in the DNA-PKcs protein (KR) abolishes its kinase activity ^22,29^ in Chinese hamster ovary cells. This increased HDR to levels 2- to 3-fold above those seen when the DNA-PKcs gene was completely inactivated ^29^.

To achieve high efficiency of targeted DSBs with few off-target DSBs we first generated an iPSC line carrying a doxycycline inducible Cas9 nickase (iCRISPR-Cas9n) as described by Gonzalez *et al.* ^30,31^. We then used this line to introduce a mutation in the *PRKDC* gene that causes the K3753R mutation in the DNA-PKcs protein ^29^. We tested the efficiency with which nucleotide substitutions can be precisely introduced in DNA-PKcs KR (KR) and wildtype DNA-PKcs cells using three substitutions that cause amino acid changes that are fixed among present-day humans but occur in the ancestral, ape-like state in the Neandertals and Denisovan genomes. These substitutions occur in the genes *CALD1*, *KATNA1* and *SLITRK1* that are involved in neurite outgrowth. Changes in them may therefore affect connectivity or neuronal circuitry in the human brain.

We designed RNA guides for double nicking ^30^ and single-stranded DNA donors carrying the mutations reverting the targeted amino acids to the ancestral states, namely V671I, A343T and A330S, in *CALD1*, *KATNA1* and *SLITRK1*, respectively. Each donor also carried silent “blocking” mutations to prevent recutting of the target once edited. When editing each gene independently, HDR efficiencies were 2.5-fold (36% edited chromosomes), 4.1-fold (79% e.c.), and 7.8-fold (51% e.c.) higher for *CALD1*, *KATNA1* and *SLITRK1*, respectively, in the KR cells than the wild type cells (Fig. 2A). This corresponds to a shift of the ratio of HDR to NHEJ from 0.23, 0.29 and 0.08 to 3.49, 25.4, and 2.60, respectively.

**Fig. 2:**
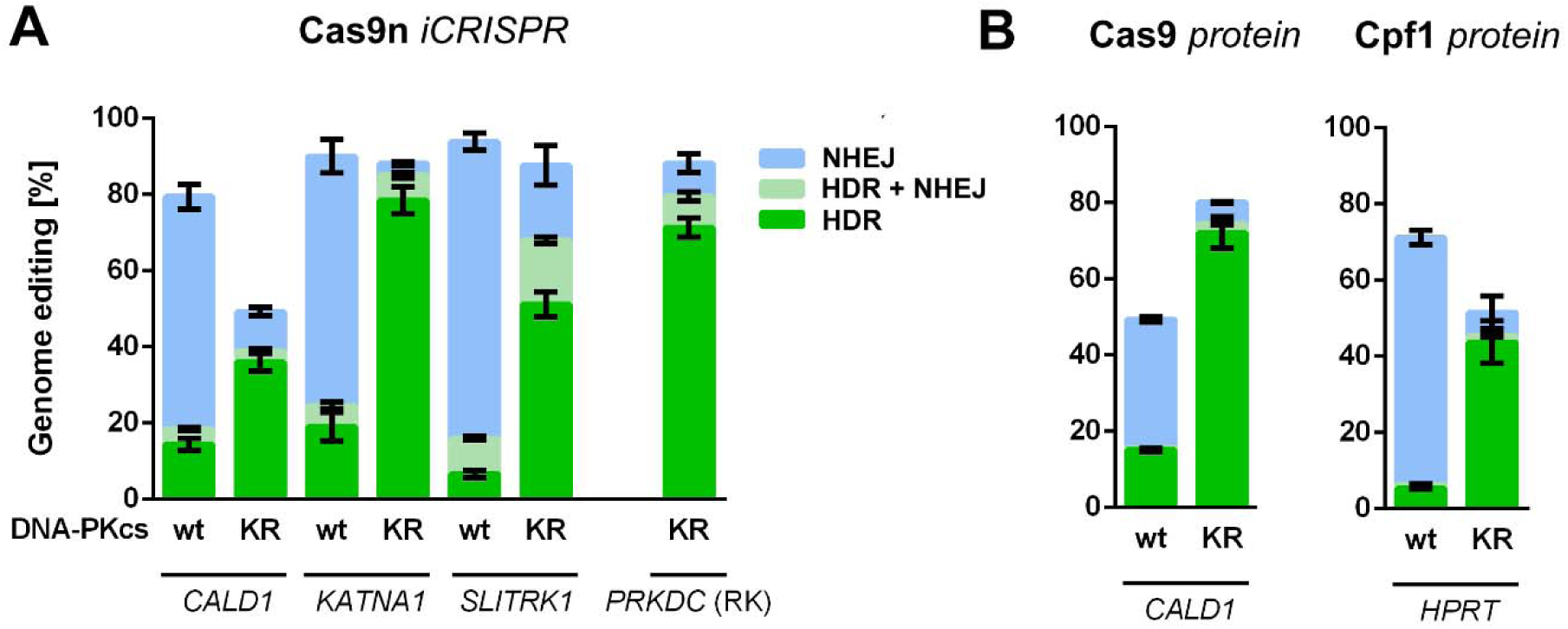
Catalytically inactive DNA-PKcs (KR) leads to increased precise genome editing. **(A)** The frequency of NHEJ and HDR in the editing of the genes *CALD1, KATNA1, SLITRK1* and *PRKDC* using Cas9n double nicking. **(B)** Editing of *CALD1* with Cas9 nuclease and *HPRT* with Cpf1 nuclease. HDR, HDR + NHEJ, and NHEJ are indicated in green, light green and blue, respectively. Error bars show standard deviation of replicate experiments.

To test if the increase in HDR relative to NHEJ was dependent on cellular production of Cas9 nickase from an endogenous inducible gene, we introduced recombinant Cas9 or Cpf1 protein, respectively, without the induction of the endogenous iCRISPR-Cas9n gene, into KR cells by electroporation and edited *CALD1* to its ancestral state and also introduced nucleotide substitutions in the *HPRT* gene in separate experiments. The precise editing of *CALD1* increased 4.8-fold and the editing of HPRT increased 8.3-fold (Fig. 2B), corresponding to a shift of the ratio of HDR/NHEJ from 0.45 to 12.8 and 0.08 to 7.1, respectively. Thus, PGE is increased in the KR cells compared to wildtype cells regardless of which enzyme is used to induce DSBs.

Encouraged by the high HDR efficiencies in the KR cells, we electroporated the six guide RNAs and the three donor DNAs for *CALD1*, *KATNA1* and *SLITRK1* into KR cells after induction of Cas9 nickase expression. When scoring HDR as the presence of any of the nine intended mutations (two silent “blocking” mutations and one amino acid replacement in each gene), we achieved HDR efficiencies of 21% for *CALD1*, 32% for *KATNA1*, and 34% for *SLITRK1* (Fig. 3A). We proceeded to isolate 33 cellular clones from the population of edited cells. Strikingly, 11 clones (33%) carry at least one targeted nucleotide substitution (TNS) on both chromosomes without any insertions or deletions in any of the three genes (Fig. 3C, left panels). Four clones (12%) carry all three ‘ancestral’ TNSs without any additional insertions or deletions, thus resulting in the three desired changes in a homozygous form (Fig. 3C, right panels).

**Fig. 3:**
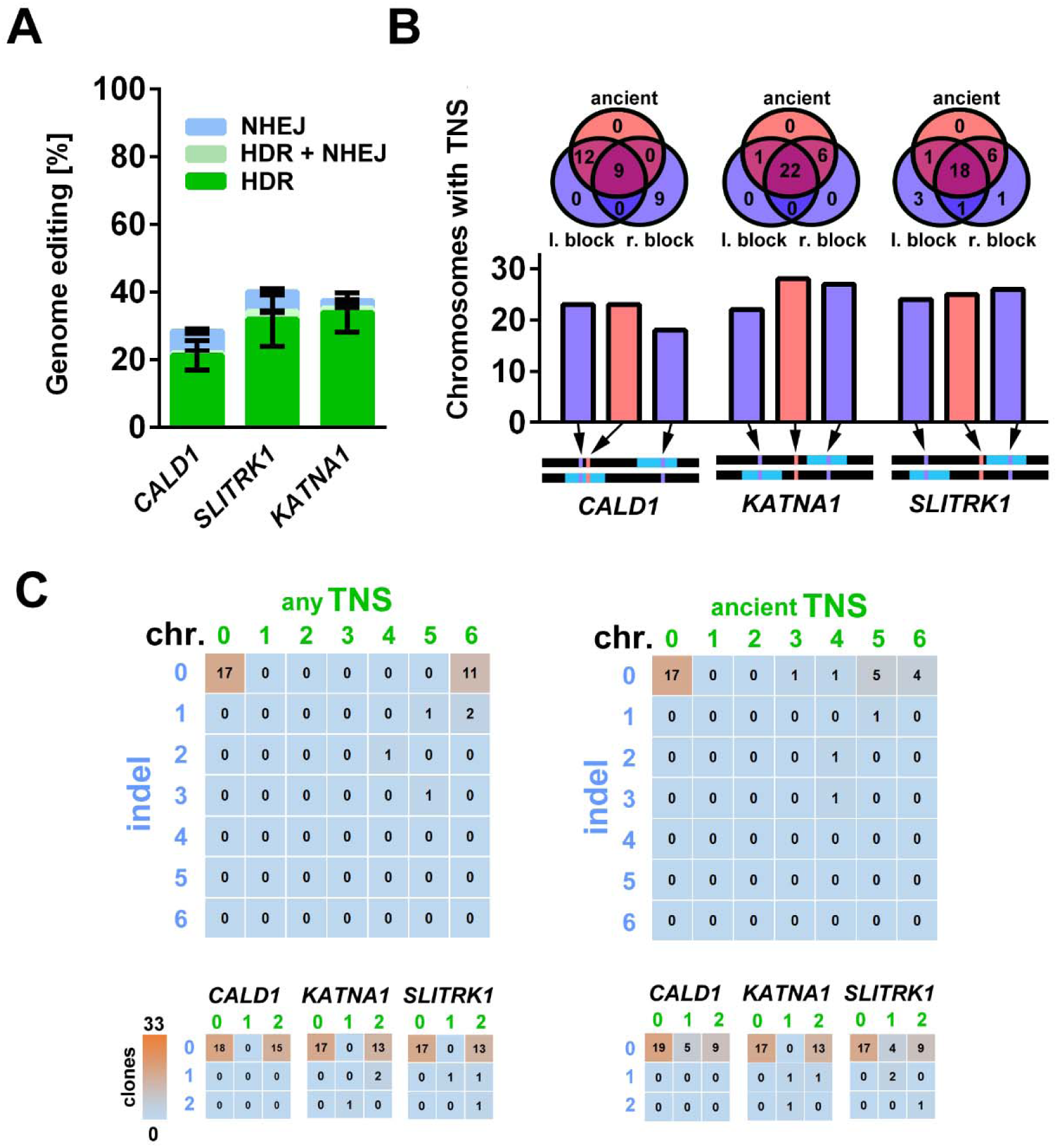
Multiplexed precise genome editing of *CALD1, KATNA1*, and *SLITRK1*. **(A)** KR cells were electroporated with the guide RNAs and DNA donors for the three genes. The frequency of HDR, HDR + NHEJ, and NHEJ for each gene is given. Error bars show the standard deviation. **(B)** Venn diagrams illustrate how often targeted nucleotide substitutions (TNS), which are the left and right blocking mutations (purple) and ‘ancient’ missense mutations (pink), were observed in the chromosomes for each gene. Blocking mutations far from the other mutations (as the right one in *CALD1*) often occur in the absence of the other mutations. **(C)** Heat maps of genome editing events (TNS and/or indels) in chromosomes carrying *CALD1, KATNA1,* and *SLITRK1* of 33 analyzed cellular clones are shown. Most clones either remain wildtype or are precisely edited for all three genes. Above all three genes (six chromosomes) with at least one TNS regardless if blocking mutation or ‘ancient’ mutation (left) or with at least the ‘ancient’ mutations (right). Below, the data for each gene (two chromosomes) is shown. An overview of mutations introduced together in *CALD1, KATNA1*, and *SLITRK1* for each of the 33 clones is in Supplementary Table 2.

The results show that the inactivation of the kinase activity of the DNA-PKcs causes almost all DSBs induced by the CRSPR nuclease in the targeted genes to be repaired by HDR rather than NHEJ (Fig. 2 and 3) indicating that NHEJ is almost completely inactivated. One concern is that this may lead to genomic instability because double-stranded DNA breaks that occur spontaneously due to both endogenous and exogenous factors the cells cannot be repaired by NHEJ. Encouragingly, the DNA-PKcs KR cells maintained a healthy karyotype with no chromosomal aberrations over 26 passages (Supplementary Fig. 1). In addition, the efficiency with which nucleotide substitutions can be introduced in these cells makes it possible to restore the normal function of the *PRKDC* gene once the desired changes have been introduced. We demonstrated this by editing the *PRKDC* gene back to its wildtype state and find that this “reverse” mutation was successfully introduced in 71% of chromosomes (Fig. 2A).

Notably, the incorporation of homozygous TNSs in all three genes is more than four-fold higher than expected by chance given the efficiencies of the introduction of substitutions when single genes are targeted. This suggests that KR cells that are “editing competent” will tend to be efficiently edited at multiple loci when presented with the relevant guide and donor nucleotides. Future work will determine the extent to which larger numbers of substitutions can be simultaneously introduced and further optimize the efficacy of multiplexed PGE.

Multiplexed PGE as described here is an advance in that it allows several substitutions to be introduced in a single round of editing. Because the lysine residue at position 3753 which we mutated in the KR cells is conserved among vertebrates (Fig. 1B) we expect that its function is also conserved so that the approach taken here is likely to be applicable not only in human but in many vertebrates. A single round of experiments as performed here where three homozygous nucleotide substitutions are introduced can be completed in two weeks. Thus, many substitutions can be introduced in a reasonable time frame. For example, it will allow the introduction of the 96 substitutions that revert 87 proteins to the primary sequences carried in the last common ancestor shared by modern humans and Neandertals ^32^. Obviously, any other combinations of substitutions of medical, evolutionary or other interest could also be generated.

## Author contributions and acknowledgements

S.R. conceived the idea and performed the experiments. S.R. and T.M and S.P. planned the experiments and wrote the paper. We would like to thank Antje Weihmann and Barbara Schellbach for DNA sequencing, and Heidrun Holland for karyotyping. This work was supported by the Max Planck Society and by the NOMIS foundation.

## Methods (in Supplementary Materials)

### Cell culture

A human iPS line (409-B2, female, Riken BioResource Center) was used to create an iCRISPR-Cas9n line as described by Gonzalez et al. ^31^(GMO permit AZ 54-8452/26). The Puro-Cas9 donor was subjected to site-directed mutagenesis with the Q5 mutagenesis kit to introduce the D10A mutation (New England Biolabs, E0554S). Mutagenesis primers were from IDT (Coralville, USA) and are shown in Supplementary Table 1. Cells were grown on Matrigel Matrix (Corning, 35248) at 37°C in a humidified incubator with 5% CO_2_ in mTeSR1 medium (StemCell Technologies, 05851) with supplement (StemCell Technologies, 05852) that was replaced daily. At ~ 80% confluency, cells were dissociated using EDTA (VWR, 437012C) and diluted 1:1 in medium supplemented with 10 µM Rho-associated protein kinase (ROCK) inhibitor Y-27632 (Calbiochem, 688000) for one day after replating.

### Design of gRNAs and single stranded DNA donors

To edit the genes *CALD1, KATNA1* and *SLITRK1* we used two gRNAs in tail-to-tail orientation with a small distance of gRNA pairs, and a cleavage efficiency above 45 (percentile rank score) as predicted by the sgRNA scorer 1.0 tool ^34^. The single stranded DNA donors contained the desired mutation close to the middle of the sequence between the nicks as well as silent blocking mutations in each gRNA target close to the PAM to prevent re-cutting. At least 30nt homology arms were upstream or downstream of the respective nick site. For Cas9 nuclease cleavage, the gRNA of the *CALD1* nickase gRNA pair that cuts closer to the desired mutation was used together with a 90nt single stranded DNA donor centered on the desired mutation and containing a blocking mutation. For Cpf1 cleavage, the positive control gRNA for *HPRT* from IDT (Coralville, USA) was used, together with a 90nt single stranded DNA donor with a blocking mutation near the PAM site and an additional mutation near the cut. RNAs and single stranded DNA donors were ordered from IDT (Coralville, USA) and are shown in Supplementary Table 1. The drop in multiplexed PGE efficiency from 33% for the introduction of at least one of the substitutions for each gene to 12% for all three missense substitutions (Fig. 3C) is mostly due to the design of the *CALD1* donor, where the right “blocking” mutation is further away from the “ancestral” mutation than in the other guide RNAs, causing this substitution to be the only one incorporated in 30% of *CALD1* TNS-positive chromosomes (Fig. 3B).#

### Lipofection of oligonucleotides

409-B2 iCRISPR-Cas9n hiPSCs were incubated in medium containing 2µg/ml doxycycline (Clontech, 631311) three days (four days for multiplexing) prior to lipofection. Lipofection was done using the alt-CRISPR protocol (IDT) at a final concentration of 7.5nM of each gRNA and 10nM of the DNA donors. In brief, 0.75µl RNAiMAX (Invitrogen, 13778075) and the respective oligonucleotides were separately diluted in 25µl OPTI-MEM (Gibco, 1985-062) and incubated at room temperature for 5min. Both dilutions were mixed to yield 50µl of OPTIMEM including RNAiMAX, gRNAs and single stranded DNA donors. The lipofection mix was incubated for 20-30min at room temperature. Cells were dissociated using EDTA for 5min and counted using the Countess Automated Cell Counter (Invitrogen). The lipofection mix, 100µl containing 25.000 dissociated cells in mTeSR1 supplemented with Y-27632, and 2µg/ml doxycycline were put in one well of a 96 well covered with Matrigel Matrix (Corning, 35248). Media was exchanged to regular mTeSR1 media after 24hours. We attempted multiplexed editing of three genes using lipofection, but achieved 3-10% HDR for a gene at best (Supplementary Fig. 2).

### Oligonucleotide and ribonucleoprotein electroporation

The recombinant *Streptococcus pyogenes* Cas9 protein, *Acidaminococcus sp. BV3L6* Cpf1 protein and respective electroporation enhancer were from IDT (Coralville, USA) and electroporation was done using the manufacturer’s protocol, except for the following alterations. Nucleofection was done using the B-16 program of the Nucleofector 2b Device (Lonza) in cuvettes for 100 µl Human Stem Cell nucleofection buffer (Lonza, VVPH-5022), containing 1 million cells, 78pmol electroporation enhancer, 160pmol of each gRNA (crRNA/tracR duplex for Cas9 and crRNA for Cpf1), 200pmol of each single stranded DNA donor, and 252pmol Cas9 or Cpf1. For multiplexing, only gRNAs and single stranded DNA donors were electroporated, since a Cas9n expressing iCRISPR-Cas9n hiPSC line was used. Cells were counted using the Countess Automated Cell Counter (Invitrogen). 90 percent of the electroporated cells were plated for bulk genotype analysis and 10 percent were plated in a separate 6well to give rise to colonies derived from a single cell (clones) for which the media was supplemented with Rho-associated protein kinase (ROCK) inhibitor Y-27632 for three days post-electroporation. After at least seven days colonies were picked for following propagation and DNA isolation.

### Illumina library preparation and sequencing

At least three days after transfection cells were dissociated using Accutase (SIGMA, A6964), pelleted, and resuspended in 15µl QuickExtract (Epicentre, QE0905T). Incubation at 65°C for 10min, 68°C for 5min and finally 98°C for 5min was performed to yield single stranded DNA as a PCR template. Primers for each targeted loci containing adapters for Illumina sequencing were from IDT (Coralville, USA) (see Supplementary Table 1). PCR was done in a T100 Thermal Cycler (Bio-Rad) using the KAPA2G Robust PCR Kit (SIGMA, KK5024) with supplied buffer B and 3 µl of cell extract in a total volume of 25µl. The thermal cycling profile of the PCR was: 95°C 3min; 34x (95° 15sec, 65°C 15sec, 72°C 15sec); 72°C 60sec. P5 and P7 Illumina adapters with sample specific indices were added in a second PCR reaction ^35^ using Phusion HF MasterMix (Thermo Scientific, F-531L) and 0.3µl of the first PCR product. The thermal cycling profile of the PCR was: 98°C 30sec; 25x (98° 10sec, 58°C 10sec, 72°C 20sec); 72°C 5min. Amplifications were verified by size separating agarose gel electrophoresis using 2% EX gels (Invitrogen, G4010-11). The indexed amplicons were purified using Solid Phase Reversible Immobilization (SPRI) beads in a 1:1 ratio of beads to PCR solution ^36^. Double-indexed libraries were sequenced on a MiSeq (Illumina) giving paired-end sequences of 2 x 150 bp (+7 bp index). After base calling using Bustard (Illumina) adapters were trimmed using leeHom^37^.

### Sequencing data analysis

Bam-files were demultiplexed and converted into .fastq files using SAMtools ^38^. CRISPresso ^39^ was used to analyse .fastq files for percentage of wildtype, HDR (any TNS), NHEJ (indels), and mix of HDR and NHEJ. Analysis was restricted to amplicons with a minimum of 70% similarity to the wildtype sequence and to a window of 20bp from each gRNA. Unexpected substitutions were ignored as sequencing putative errors. Sequencing data from colonies derived from single cell seeding (Fig. 3B and C) was analyzed using SAMtools and colonies were regarded as clones if the clear majority of reads consisted of a single sequence (homozygous) or of two sequences of similar read count (heterozygous) and blocking mutations, ancient mutations, as well as indels for each chromosome of the cells of a clone were noted (Supplementary Table 2).

### Karyotyping

Microscopic analysis of the karyotype of 25 metaphases was done after trypsin induced Giemsa staining. The analysis was carried out in line with international quality guidelines (ISCN 2016: An International System for Human Cytogenetic Nomenclature ^40^) by the ‘Sächsischer Inkubator für klinische Translation’ (Leipzig, Germany).

## Supplementary Data

**Supplementary Fig. 1:**
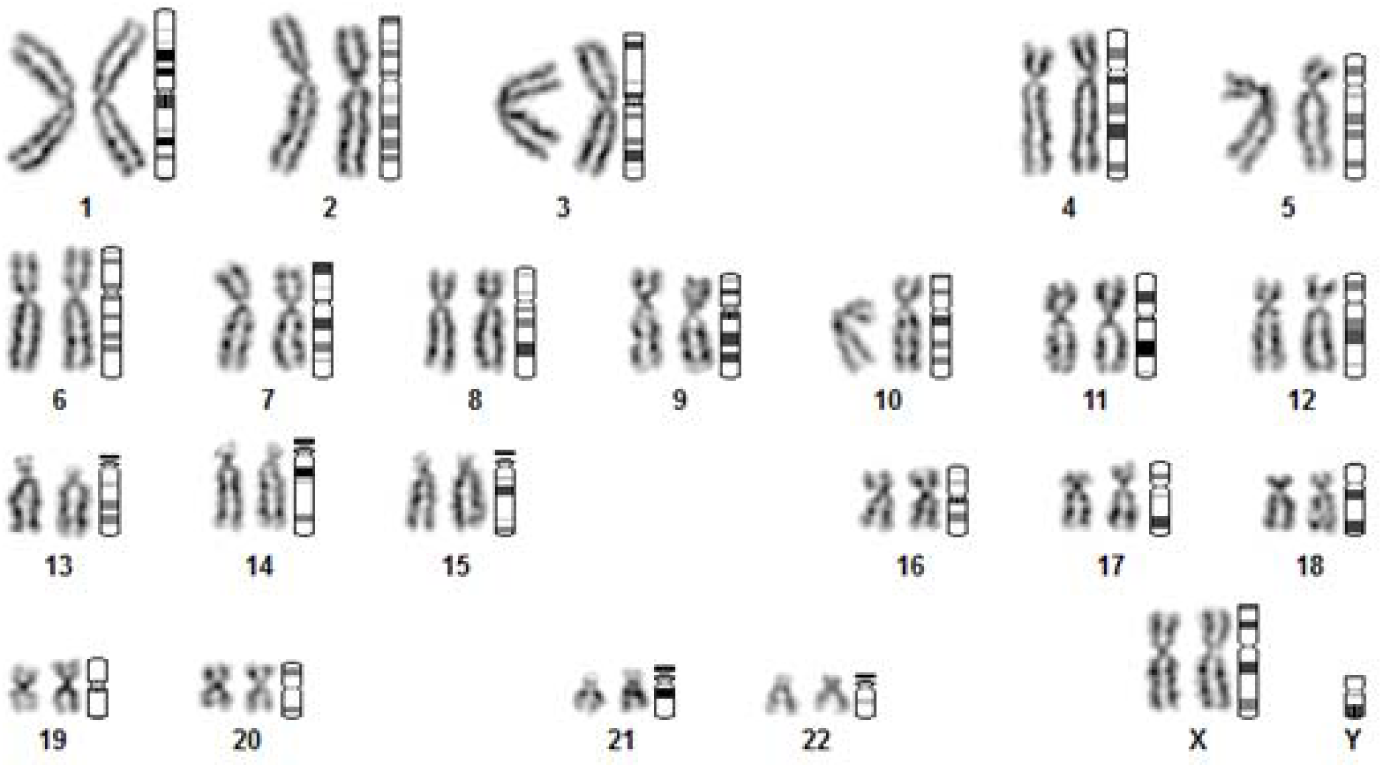
Representative karyogram of 409-B2 iCRISPR hiPSCs with the DNA-PKcs KR mutation after 3 months in culture. Of 25 metaphases, analyzed by trypsin induced Giemsa staining, all show a healthy karyotype (46, XX). No numerical or large scale chromosomal aberrations were identified (band number 350, gray shades 3).

**Supplementary Fig. 2:**
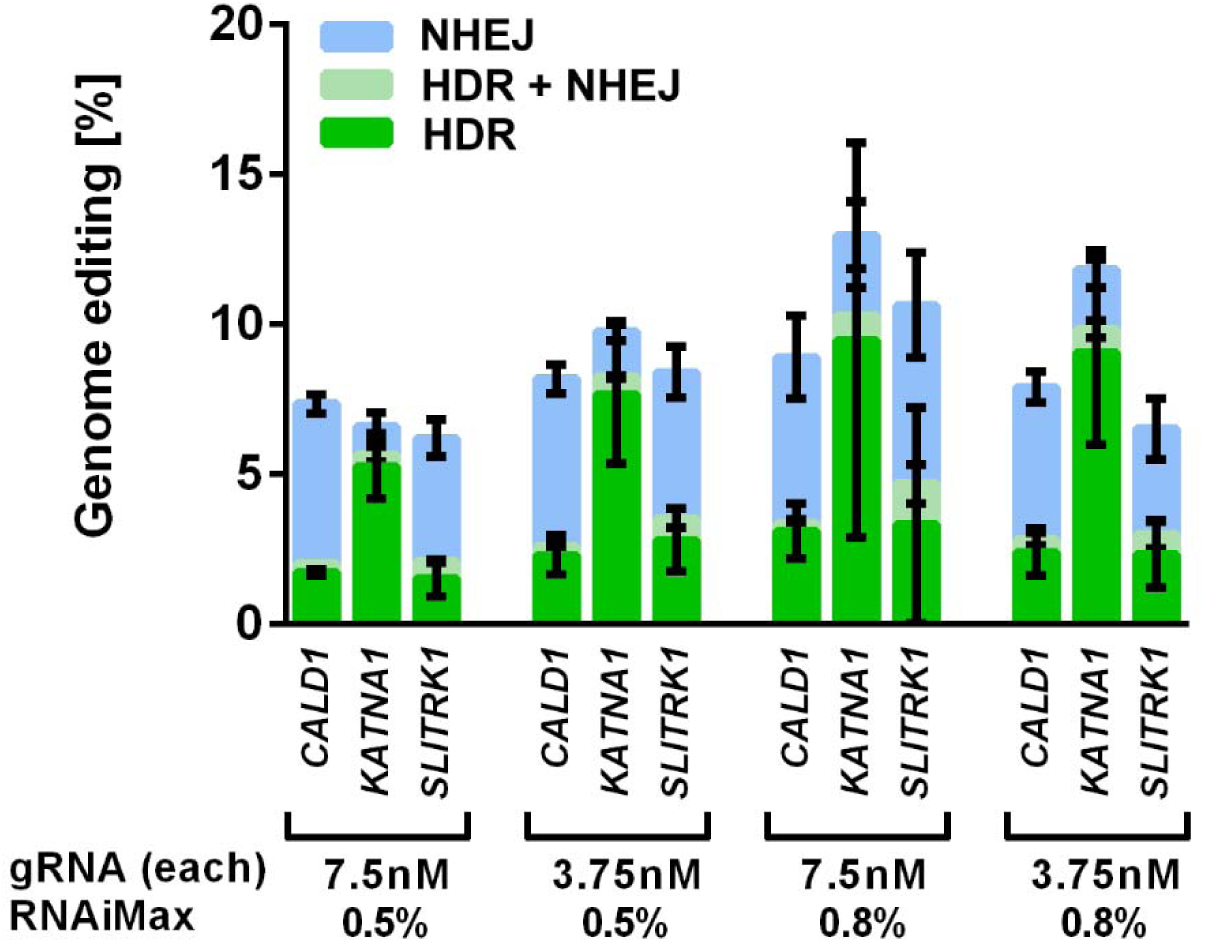
Genome editing efficiencies after multiplexed delivery of gRNAs and single stranded DNA donors of *CALD1, KATNA1*, and *SLITRK1* by lipofection. Lipofection of gRNAs and single stranded DNA donors allows only low bulk HDR efficiencies for all three genes in 409-B2 hiPSCs with the DNA-PKcs KR mutation. Combinations of two gRNA and two lipofection reagent (RNAiMax) concentrations were tested. HDR, HDR + NHEJ, and NHEJ are shown in green, light green or blue, respectively. Error bars show the standard deviation of three technical replicates and cells were incubated for four days with doxycycline to express Cas9n for double nicking.

**Supplementary Table 1:**
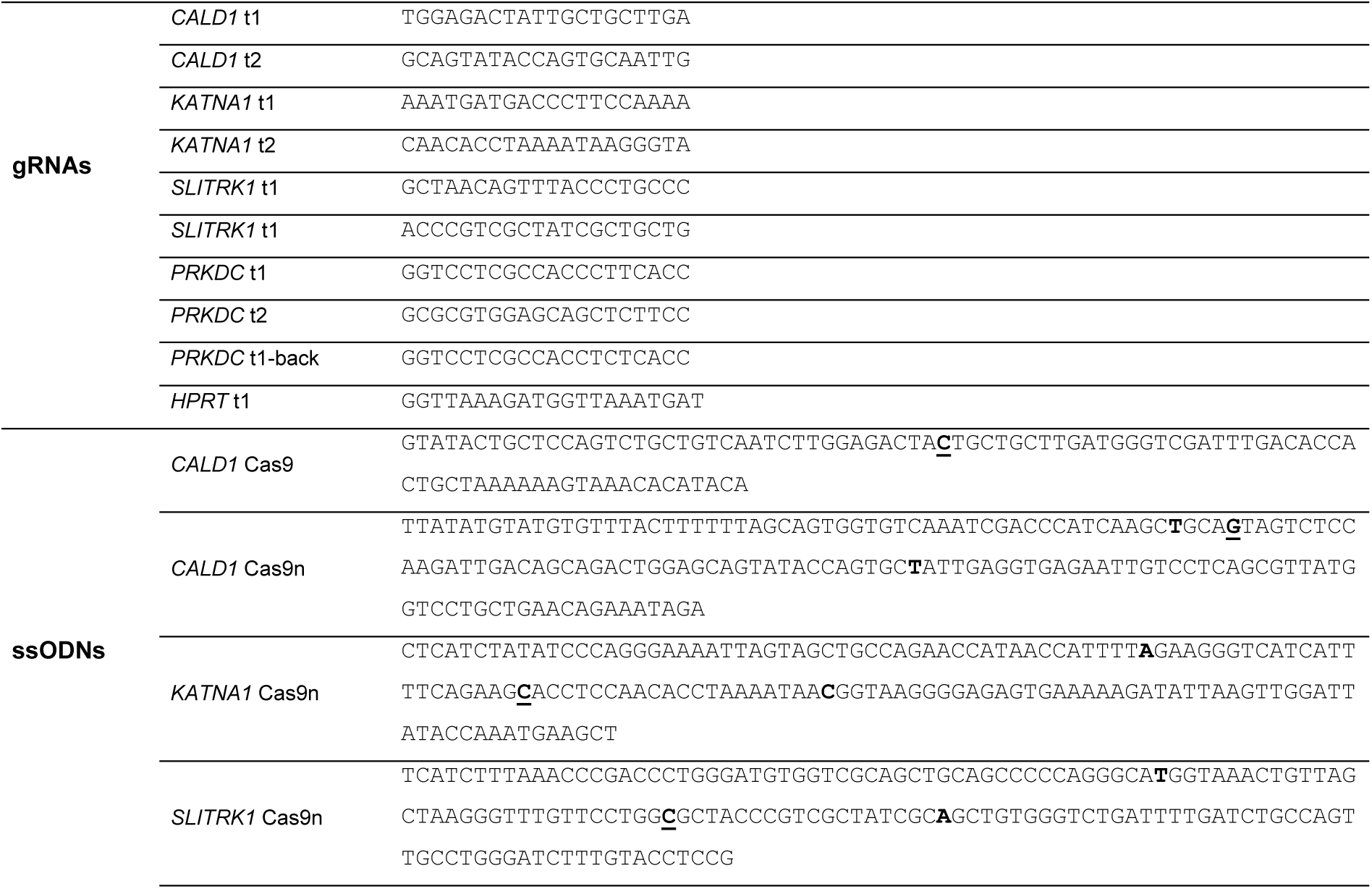
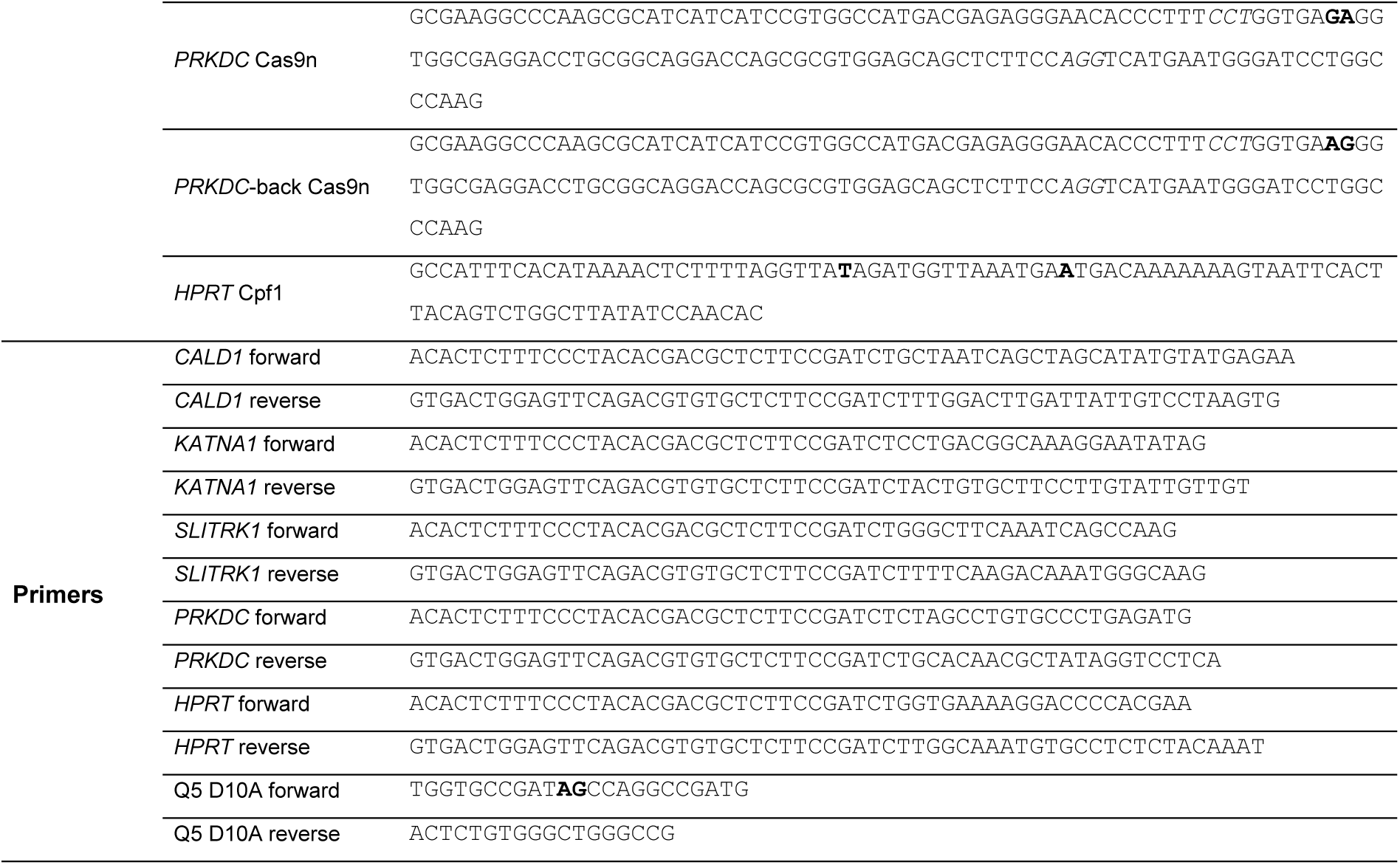
Oligonucleotides used. gRNA (crRNA target) and single stranded DNA donors (ssODNs) for editing of *CALD1*, *KATNA1*, *SLITRK1*, *PRKDC*, and *HPRT*, as well as primers for analysis and Q5 site-directed-mutagenesis of the Cas9 iCRISPR donor plasmid are shown. Mutations are in bold letters and ancestral mutations are underlined as well. The gRNA ‘*CALD1* t1’ was used for Cas9n and Cas9 cleavage.

**Supplementary Table 2:**
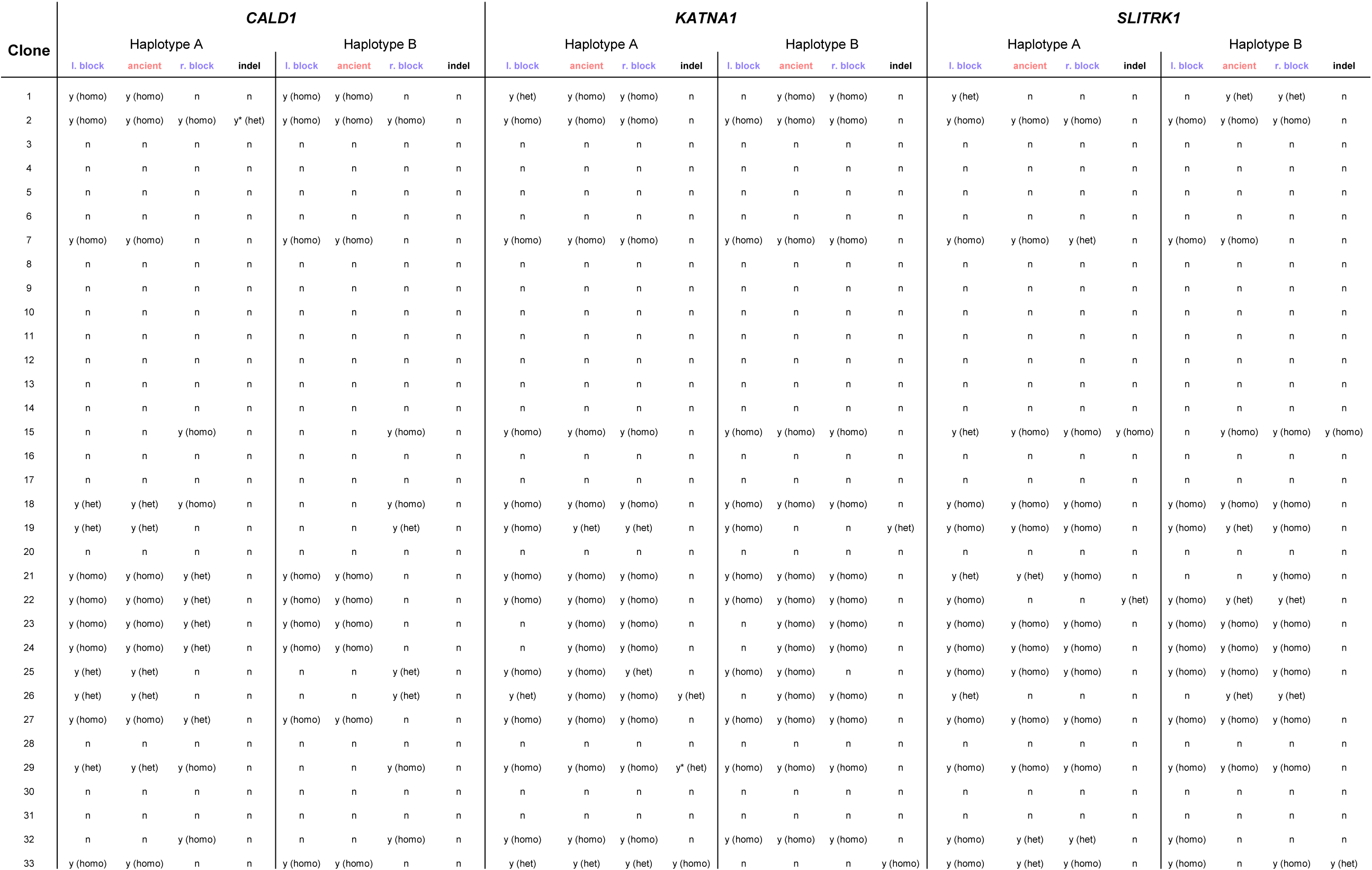
Overview of mutations introduced in *CALD1, KATNA1*, and *SLITRK1* in 33 cellular clones. Integration of targeted nucleotide substitutions (left blocking mutation, ‘ancient’ missense mutation, and right blocking mutation) and insertion/deletions (indels) is labelled with ‘y’, while absence of these mutations is labelled with ‘n’. Homozygous (homo) or heterozygous (het) integration of the mutations is stated. ‘y*’ indicates two cases where non-targeted missense nucleotide substitutions occur (clone 2 in *CALD1* and clone 29 in *KATNA1*).

